## Supplementary figures and images for "Genetic and compound screens uncover factors modulating cancer cell response to indisulam"

### Supplemental Figures

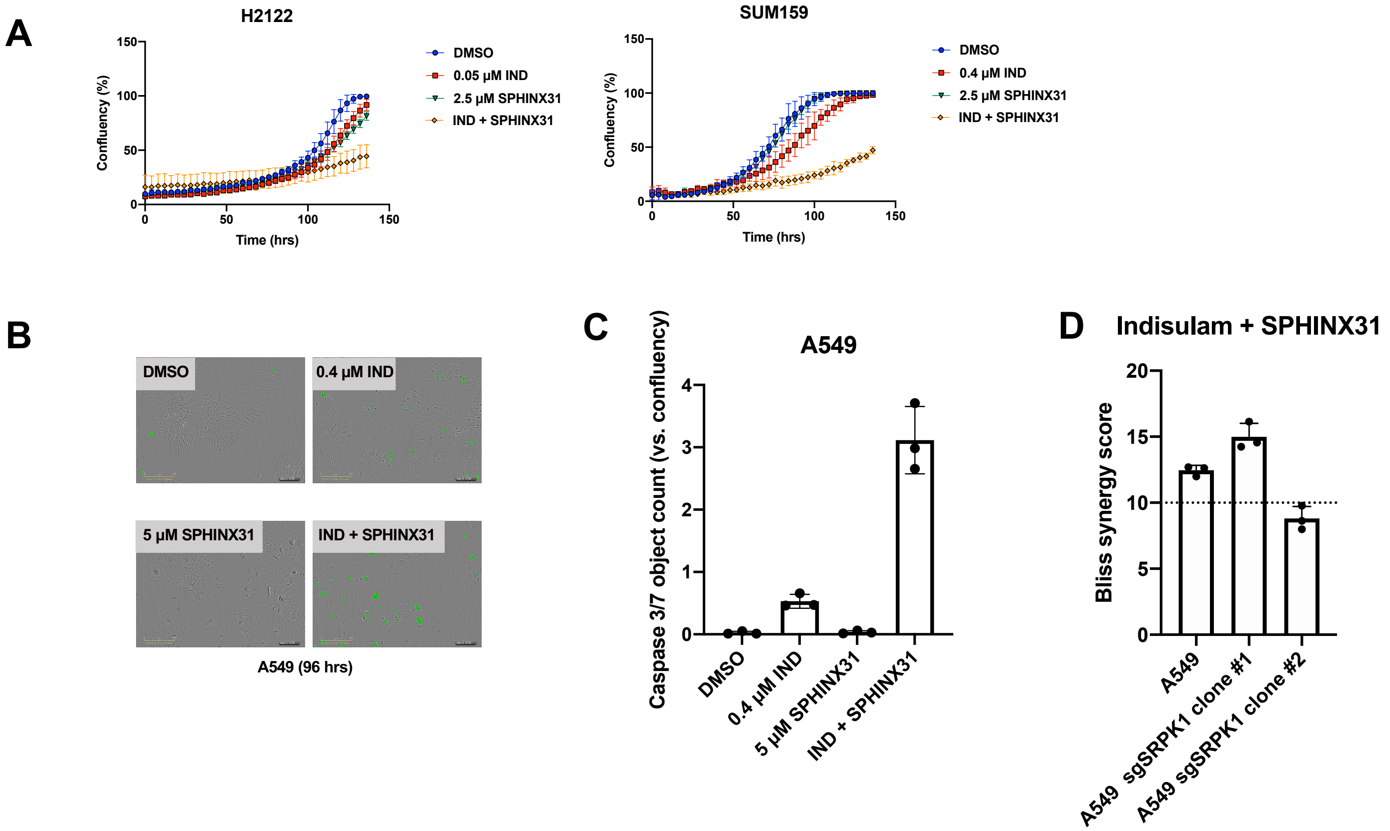


Supplemental Figure 1

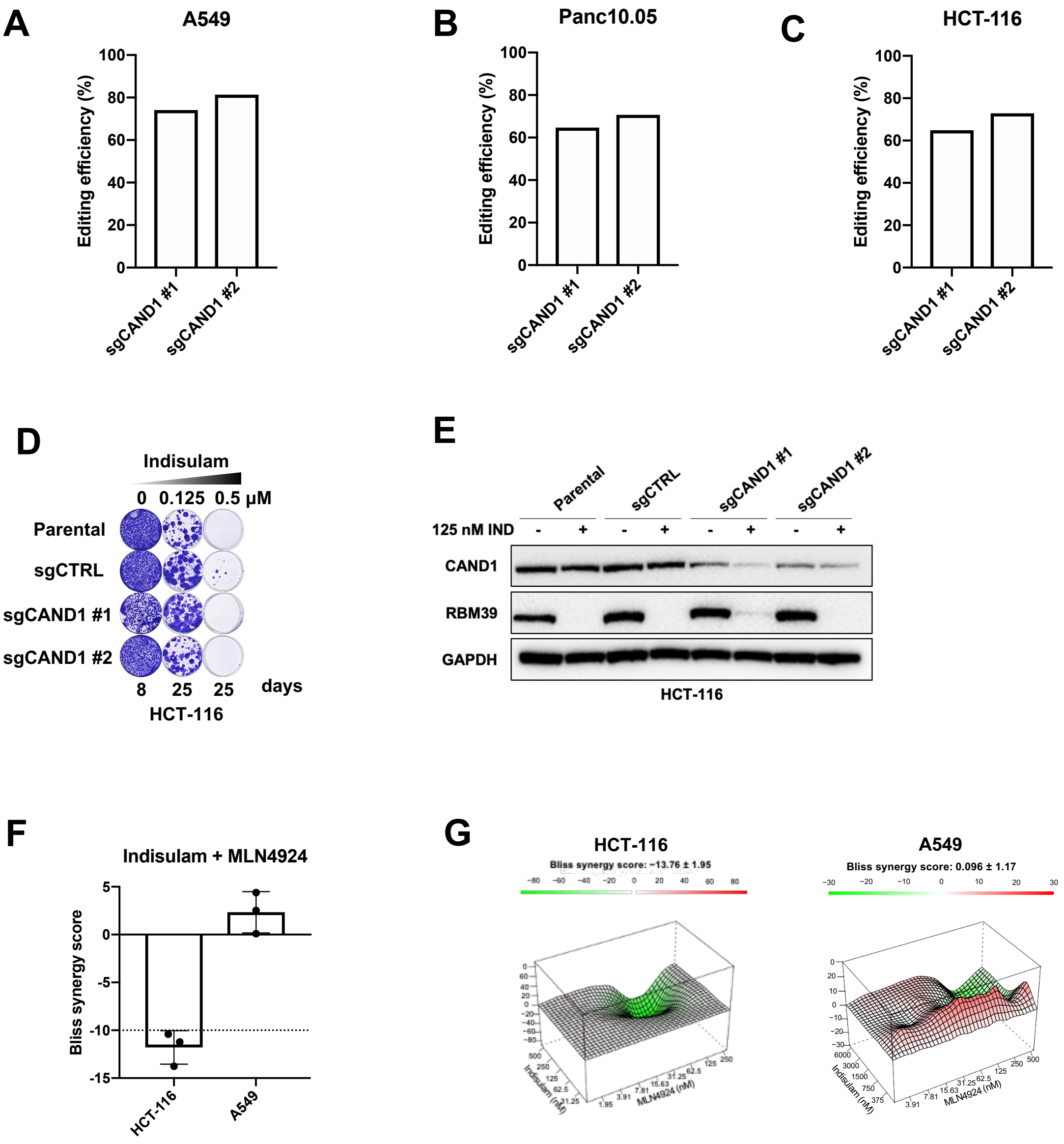


Supplemental Figure 2

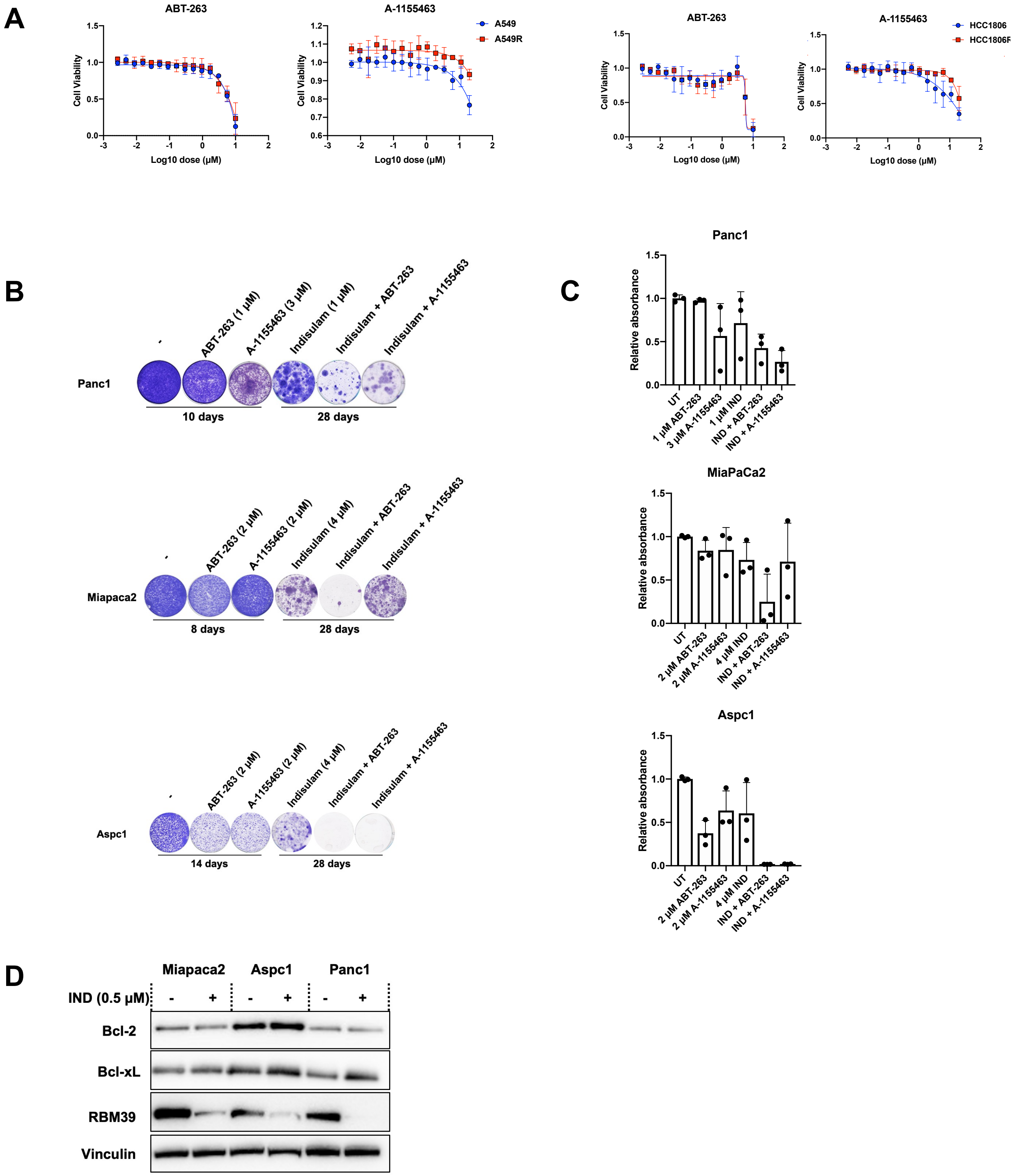


Supplemental Figure 3
